# Lipid anchor engineering controls cell-penetrating arginine-rich peptide presentation for efficient siEGFR liposomal delivery to triple-negative breast cancer cells

**DOI:** 10.64898/2026.08.20.745705

**Authors:** Piotr Białecki, Simone Braccia, Tomasz Makowski, Kinga Piórecka, Lucia Falcigno, Rosa Bellavita, Annarita Falanga, Maria Bryszewska, Agnieszka Robaszkiewicz, Stefania Galdiero, Elżbieta Pędziwiatr-Werbicka

## Abstract

Understanding the physicochemical factors that govern siRNA nanocarrier assembly is essential for the rational design of effective delivery systems. By optimizing various lipid compositions, cholesterol content and PEG length we created a peptide-functionalized cationic liposomal platform made of DOPE/TAP lipids with cholesterol-anchored nona-arginine (R9-Chol) for siRNA complexation, intracellular transport and effective silencing of the target *EGFR* gene. Analysis of ζ-potential and dynamic light scattering allowed to rationally design formulation of stable, monodisperse nanoscale lipoplexes with a positive surface charge. With fluorescence polarization, circular dichroism and agarose gel electrophoresis we found an optimal siRNA:liposome complexation ratio of 1:77, which protected siRNA from ribonuclease-mediated degradation. Morphological imaging confirmed a shift from discrete vesicular structures to organized multilamellar lipoplexes, consistent with electrostatically driven self-assembly. In cellular studies, the optimized nanocarrier promoted efficient uptake of fluorescent siRNA in MDA-MB-231 cells and achieved functional delivery of anti-*EGFR*, leading to substantially reduced expression of the target gene at both transcript and protein levels. This work offers mechanistic understanding of peptide-assisted lipid–siRNA assembly and positions R9-functionalized DOPE/TAP liposomes as a promising platform for siRNA delivery.

## Introduction

Liposomes are among the most widely studied nanoparticle-based drug delivery systems because of their biocompatibility, structural flexibility, and ability to encapsulate a wide range of therapeutic agents, including nucleic acids, proteins, and small molecules [1][2][3][4]. Their amphiphilic phospholipid structure allows them to self-assemble into bilayer vesicles in aqueous environments, creating a biomimetic platform for controlled transport of cargo. Notably, liposomal systems can be easily modified through surface functionalization to improve cellular uptake, stability, and delivery efficiency.

Cell-penetrating compunds such as cationic arginine-rich peptides (CARPs) have been known for their intrinsic neuroprotective properties [5], but also as useful tools for enhancing intracellular delivery of therapeutic nucleic acids. Among these, nona-arginine (R9), a guanidinium-rich cationic peptide, shows strong membrane translocation ability by forming bidentate hydrogen bonds with anionic cell surface components (phosphate and sulfate), initiating cellular uptake, as well as high affinity for negatively charged biomolecules through electrostatic interactions. These features make R9 an appealing functional component for plasmid DNA and siRNA delivery systems [6][7]. The structural properties of CARPs allow their various chemical modifications for coupling with other molecules or nano-platforms, which were recently reviewed by Anjous R and co-authors, to facilitate their intracellular delivery. In former attempts nona-arginine (R9) peptides were enzymatically linked to, for example, anti-Egfr nanobodies, conjugated to a helical template designed with a repeating sequence l-Leu-l-Leu-Aib, a small hydrophobic peptide anchors and to fatty acids [8], or subjected to peptidomimetic conversion [9].

Bearing in mind the advantage of the liposome properties and cationic arginine-rich peptides, we hypothesize that the nona-arginine (R9) peptide can be incorporated into liposome lipid bilayer, thereby creating liposome-R9 peptide platform for efficient delivery of siRNA. The basal composition of the liposome was selected to ensure cationic properties of at least one phospholipid, which aimed to avoid the electrostatic interaction of the peptide with the liposome structure. As a cargo attachment into lipid bilayer, we tested DSPE-PEG2000 maleimide as a standard amphiphilic lipid anchor for peptide display as well as cholesterol as a membrane-integrating anchor linked through a short PEG spacer to improve peptide accessibility and maintain liposomal stability [10]. Moreover, we used various lipid:cholesterol:PEG:R9 ratios to find the optimal composition for the interaction with siRNA.

To validate the efficacy of intracellular siRNA delivery we targeted epidermal growth factor receptor (Egfr), a receptor tyrosine kinase often dysregulated in aggressive tumors, including triple-negative breast cancer (TNBC), where it drives proliferation, migration, invasion, and therapy resistance [11]. The chosen MDA-MB-231 cell line is characterized by moderate *EGFR* expression to eliminate the risk of Egfr high-dose paradox [12]. Anti-Egfr approaches are considered as monotherapy or in combination with chemotherapy drugs for the treatment of TNBC, but the ongoing clinical trials have been testing Egfr inhibitors and anti-Egfr antibodies only. Despite promising outcomes in the *in vitro* and mouse models, siRNA delivery faces several challenges such as poor biodistribution or crossing cell membrane, susceptibility to lysosomal and enzymatic degradation. At least some of these challenges including uptake of liposomes by the mononuclear phagocyte system or endonuclease cleavage in the blood stream can be prevented by PEGylation, which improves colloidal stability and prolongs circulation time, as well as by surface-displayed cationic peptides, which enables efficient surface loading, facilitate intracellular uptake and enhance endosomal escape through membrane destabilization.

In this study we designed, developed and validated the new R9-modified liposomes capable of efficient si*EGFR* delivery into cells, which was followed by biologically meaningful knockdown of the receptor in the considered cells.

## 2. Materials and methods

### 2.1 Materials

Amino acids amino acids, Fmoc-Arg(Pbf)-OH, Fmoc-Cys(Trt)-OH were acquired from GL Biochem Ltd. (Shanghai, China). N,N′-diisopropylcarbodiimide (DIC), Oxyma pure, 1-[Bis(dimethylamino)methylene]-1H-1,2,3-triazolo[4,5-b]pyridinium 3-oxid hexafluorophosphate (HATU), N,N-Diisopropylethylamine, triisopropylsilane (TIS), 1,1,1,3,3,3-Esafluoropropan-2-olo (HFIP) were purchased from Sigma-Aldrich (Italy). Rink amide p-methylbenzhydrylamine (MBHA) resin, Fmoc-L-Lys(Mtt)-OH, Fmoc-O2Oc-OH, piperidine and trifluoroacetic acid (TFA) were purchased from Iris Biotech GmbH (Marktredwitz, Germany). Solvents N,N-Diisopropylethylamine (DIPEA), N,N-dimethylformamide (DMF), Dichloromethane (DCM), Dimethyl sulfoxide (DMSO), Chloroform, Ethanol and Cholesteryl Chloroformate were purchased from Sigma-Aldrich (Milan, Italy). 1,2-di-(9Z-octadecenoyl)-sn-glycero-3-phosphoethanolamine (DOPE); 1,2-dimyristoyl-3-trimethylammonium-propane (TAP), 1,2-distearoyl-sn-glycero-3-phosphoethanolamine-N-[maleimide(polyethylene glycol)-2000] (DSPE-PEG(2000) Maleimide) were purchased from Avanti Polar Lipids, Inc. (Alabaster, AL, USA). Lipofectamine RNAiMAX, OptiMem, SuperSignal™ West Pico Chemiluminescent Substrate, PageRuler™ Prestained Protein Ladder (#01154870), Pierce™ Protease Inhibitor Tablets (EDTA-free; PIC), CellMask™ Plasma Membrane Stains Deep Red, SlowFade™ Glass Soft-set Antifade Mountant (with DAPI), RNase A, High-Capacity cDNA Reverse Transcription Kit, TRI Reagent, PowerUp SYBR Green Master Mix and oligonucleotides for real-time PCR were from Thermofisher Scientific (Thermofisher Scientific, Warsaw, Poland). Heparin sodium salt from porcine intestinal mucosa was purchased from Sigma-Aldrich (Merck, Poznan, Poland). Anti-Egfr rabbit mAb (#4267), Histone H3 Rabbit Ab (#9715), anti-rabbit IgG, HRP-linked Antibody (#7074) were from Cell Signaling Technologies (LabJOT, Warsaw, Poland). Agarose was from Maximus (Maximus, Lodz, Poland) and GelRed stain was purchased from Biotium, Inc., (Fremont, CA, USA). All chemicals and reagents were of analytical grade and used as received.

### 2.2 siRNA

Liposomes DOPE/TAP/R9-Chol were complexed with non-fluorescent or FITC-labelled siRNA silencing *EGFR* (sense: 5’ G.C.U.A.U.G.A.G.A.U.G.G.A.G.G.A.A.G.A.U.U 3’; antisense: 5’-P U.C.U.U.C.C.U.C.C.A.U.C.U.C.A.U.A.G.C.U.U 3’). SiRNA was purchased from Dharmacon Inc., Lafayette, CO, USA. siRNA was dissolved in 1xsiRNA buffer (Dharmacon Inc., Lafayette, CO, USA).

### 2.3 Cell culture

MDA-MB-231 cells were cultured in DMEM supplemented with 10% FBS, penicillin–streptomycin (50 U/mL and 50 μg/mL, respectively) in 5% CO_2_. Culture medium was replaced 2x a week.

### 2.4 Peptide synthesis and purification

Two different strategies were used for the inclusion of R9 in the liposome formulation in order to have the R9 located on the surface of liposomes [13]. The first strategy involved the binding of R9 to DSPE-PEG while the second strategy was the binding to cholesterol.

The peptides were synthesized using the Rink amide resin (0.67 loading) as solid support. The Fmoc protecting group was removed by treatment with a solution of 30% piperidine in DMF (2 × 15 min). Fmoc-Lys(Mtt)-OH was used as the first amino acid performing the conjugation of the cholesterol on the amine group in the lysine side chain; while a Cys residue was introduced at the C-terminus as the first amino acid in the conjugation to DSPE-PEG. Each reaction coupling was performed through 2 coupling steps. In the first, Fmoc-amino acid (2 eq) was added with N,N′-diisopropylcarbodiimide (DIC, 2 eq) oxyma pure (2 eq) as coupling reagents, in DMF for 40 min; in the second, Fmoc-amino acid (2 eq) was added with HATU (2 eq), DIPEA (4 eq), in DMF for 40 min.

In the first strategy, the cysteine residue at the C-terminus of R9 was used for binding to DSPE-PEG(2000) maleimide. The peptide was cleaved from the resin with an acid solution of TFA/H_2_O/TIS (92.5/2.5/5, v/v/v) under agitation overnight. Then the peptide was precipitated in diethyl ether and separated by centrifugation (2 x 15 min, 6000 rpm). The crude peptide was dissolved in HFIP (10%) and H2O (0.1 % TFA) and purified by RP-HPLC semi-preparative (Jasco) on a Luna Omega 5 μm Polar C 150 x 21.2 mm column, with a linear gradient of solvent B (0.1% TFA in acetonitrile) in solvent A (0.1% TFA in water) from 50 % to 95 % for 15 min. DSPE-PEG(2000) maleimide was dissolved in DMF 1mg/mL and added dropwise using the syringe pump flow rate 25 μL/min to a solution of R9 under magnetic stirring (500 rev/min) at 40 °C. Then the solvent was removed using a rotavapor, freeze-dried and the R9-DSPE-PEG2000 reaction purified using the semi-preparative HPLC (Jasco) on a Jupiter 10u C18 300A 250 x 21.2 mm column with a linear gradient of solvent B (0.1% TFA in acetonitrile) in solvent A (0.1% TFA in water) from 10 % to 90 % for 20 min.

For the second strategy, the Mtt group of the lysine was removed, covalently conjugating the cholesterol on the side chain. The Mtt deprotection was performed by treating the resin with a cocktail of DCM:TFA:TIS (94:1:5, *v*/*v*/*v*). Once complete, the Mtt removal was ascertained by the colorimetric Kaiser test used for the detection of primary amines in the solid phase. Cholesterol was added using cholesteryl chloroformate (2 eq), DIPEA (4 eq) in DMSO (2 x 2h). The peptide was then cleaved from the resin with an acid solution of TFA/H_2_O/TIS (92.5/2.5/5, *v*/*v*/*v*) under agitation overnight. The peptides were then precipitated in diethyl ether and separated by centrifugation (2 x 15 min, 6000 rpm). The crude peptide was dissolved in HFIP (10%) and H_2_O (0.1 % TFA) and purified by RP-HPLC semi-preparative (Jasco) on a Luna Omega 5 μm Polar C 150 x 21.2 mm column, with a linear gradient of solvent B (0.1% TFA in acetonitrile) in solvent A (0.1% TFA in water) from 50 % to 95 % in 15 min.

### 2.5 NMR experimental section

NMR analyses were carried out in DMSO-d6 (99.8% D, Sigma Aldrich) for R9_C-mal-DSPE-PEG and in CD_3_OH (99.8% D, Sigma Aldrich) at 298 K and using 600 MHz or 700 MHz Bruker Advance spectrometers located at the Department of Pharmacy – University “Federico II” of Naples and equipped with a z-gradient 5mm triple-resonance cryoprobe. Sample solutions was prepared by dissolving weighted amounts of products in 600-750 µL of deuterated solvents. 1D-NMR spectra were recorded with a preparation delay d1 time of 1s and of 30s and analyzed using MESTRENOVA 6.0 software (Mestrelab Research, S.L, Santiago de Compostela, Spain). 2D homonuclear spectra such as TOCSY (mixing time 70 ms) was recorded in the phase-sensitive mode using the method from States, using 4096 data points in t2 and 512 equidistant t1 values and analyzed using the CARA program (http://cara.nmr.ch/doku.php/home). Chemical shifts were referenced to 2.49 ppm, as residual CHD_2_ proton of DMSO and to 3.35 ppm, as residual CHD_2_ proton of deuterated methanol.

### 2.6 Fourier-transform infrared spectroscopy (FTIR) analysis

R9 peptide, cholesterol, and the R9-Chol construct were analysed by FTIR, using a Jasco FT/IR-4X spectrometer equipped with a DLaTGS detector. Spectra were recorded in transmittance mode over the frequency range of 400-4000 cm⁻¹.

### 2.7 Liposome preparation

Liposomes were prepared at a lipid concentration of 0.1 mM according to the extrusion method[14],[15]. In particular, lipids were dissolved in chloroform and according to the strategy, R9-CHOL in ethanol or R9-DSPE-PEG in chloroform was added; the solvents were then evaporated under nitrogen stream to obtain lipid. Lipid films were hydrated with PBS 1X for 1h freeze-thawed 6x and extruded 10x through a polycarbonate membrane with 0.1 μm diameter pores to obtain large unilamellar vesicles (LUVs).

### 2.8 Preparation of siRNA:liposome complexes

siRNA:liposome complexes were prepared by mixing the liposomal formulation containing the peptide R9 to siRNA to produce the siRNA:liposome complex. The siRNA:liposome ratio is always reported relative to the total amount (in moles) of lipids present in the formulation. siRNA solution was added at room temperature in a 2 mL eppendorf with an incubation time of 15 min. The liposome formulation was used at different concentrations for different ratios of the complex.

### 2.9 The physical characteristics of the liposome/siRNA complexes

The liposome/siRNA:liposome complexes size and polydispersity index (PDI) were measured by dynamic light scattering (DLS) using a Zetasizer NanoZS (Malvern, UK). Particle size and PDI were performed in nuclease-free water or phosphate buffer 10 mM at 25°C by calculating the average of 3 measurements.

including their hydrodynamic diameter and polydispersity index (PDI), were assessed via dynamic light scattering (DLS) using a Zetasizer NanoZS (Malvern, UK). These measurements, along with the potential analysis, were conducted at 25°C in either nuclease-free water or a 10 mM phosphate buffer, with results representing the average of three independent runs. To investigate the stability and formation of the complexes, fluorescence polarization was employed using FITC-labeled siRNA and a PerkinElmer LS-50B spectrofluorometer. The fluorescence polarization was measured according to our protocol published in [16]. Time stability of complexes formed in complete binding ratio (siRNA/liposome) were conducted every 15 min for the first 240 min (4 h). Additionally, gel electrophoresis was used to determine the complexation ratio and evaluate the complexes’ stability against ribonucleases according to the protocol [17]. Results were captured via a Gel Imaginer Azure (Azure Biosystems, Dublin, CA).

Structural insights into nucleic acid changes were obtained through circular dichroism (CD) using a Jasco J-815 spectrometer (Jasco International Co., Ltd., Tokyo, Japan) according to the protocol [18]. Baseline and probes were measured in phosphate buffer (10 mmol/L, pH = 7,4) at a wavelength set from 200 to 320 nm. The morphology of the particles and complexes was visualized using transmission electron microscopy (TEM) on a JEOL-1010 instrument (JEOL,Tokyo, Japan). Sample solutions (10 µl) were put on 200-mesh copper grids with a carbon surface and stained with uranyl acetate solution for 3 min. The grids were washed with deionized water and finally dried at room temperature.

### 2.10 Gel electrophoresis

The determination of the liposome complexation ratio with si*EGFR* was performed according to the protocol [16]. Gel electrophoresis was performed to assess the stability of the complex in the presence of ribonucleases according to the protocol described in [17]. RNAase A was added to each sample at a concentration of 1.25 μg/ml. Samples were incubated at 37 °C for 2 h. After this time a heparin diluent of 0.082 mg/ml was added to one of the 2 samples in each sample concentration and incubated on ice for 15 min. The separation process was carried out at a voltage of 90 V and a current of 35 mA for 45 min. Visualization was accomplished using ultraviolet (UV) light, and a digital image of the stained gel was captured with a Gel Imaginer Azure (Azure Biosystems, Dublin, CA).

### 2.11 Atomic forces microscopy (AFM) analyses

The morphological analysis of DOPE/TAP/R9-Chol was conducted using atomic force microscopy (AFM) according to the protocol [18]. Image processing was performed using the Scanning Probe Image Processor Software (SPIP) by Image Metrology, Hørsholm, Denmark.

### 2.12 Confocal microscopy

Accumulation of siRNA:liposome complex was performed by monitoring of the fluorescent siRNA according to our protocol [19]. Complex accumulation was measured after 3 h incubation. si*EGFR* was labelled with fluorescein (siRNA-FITC). Cell membranes were stained with CellMask™ Plasma Membrane Stains. The following wavelength values of excitation and emission were used for specimen visualization: 405 and 430–480 nm for DAPI, 649 and 666 nm for CellMask and 498 and 517 nm for siRNA-FITC.

### 2.13 siRNA accumulation analysis by flow cytometry

The quantitative level of uptake of the tested liposomes was assessed by measuring si*EGFR*-FITC using flow cytometry according to [20] The fluorescence intensity was measured by a flow cytometer (LSR® II Becton Dickinson) at excitation 470 nm and emission 595 nm for fluoresceine, The distribution of cell intensity was analyzed using FlowJo™v10.8 Software (BD Life Sciences; RRID:SCR_008520). The analyzed cell population was discriminated based on FSC-A and SSC-A parameters. The intensity of cell fluorescence was visualized on a histogram and the shift in fluorescence distribution indicated the alteration in siRNA. For quantification, the average of fluorescence intensity in every sample was used.

### 2.14 Western Blot

The level of effectiveness of inhibition of protein expression by the tested siRNA:liposome complexes was determined according to the protocol described in [21]. Proteins were separated by SDS–PAGE, transferred into a nitrocellulose membrane, and stained with primary antibodies (1:5000) at 4°C overnight. After subsequent staining with HRP-conjugated secondary antibodies (1:5000 for antirabbit and 1:2500 for anti-mouse antibodies; room temperature; 2 h), The signal was developed with the SuperSignal™ West Pico Chemiluminescent Substrate and pictures were acquired using ChemiDoc-IT2 (UVP, Meranco, Poznan, Poland). H3 was used as the loading control.

### 2.15 Real time – PCR

mRNA expression levels were determined by first extracting total RNA with TRI Reagent™ (according to [22]) and then performing reverse transcription using the High-Capacity cDNA Reverse Transcription Kit. Real-time PCR analysis was conducted on a Bio-Rad CFX96 C1000 Touch system, utilizing PowerTrack™ SYBR™ Green Master Mix and specific primers designed for this study. As per standard protocol, gene expression data were normalized to internal housekeeping controlsThe ratio between the studied and housekeeping genes was assumed to be 1 for control cells.

**Table 2.**
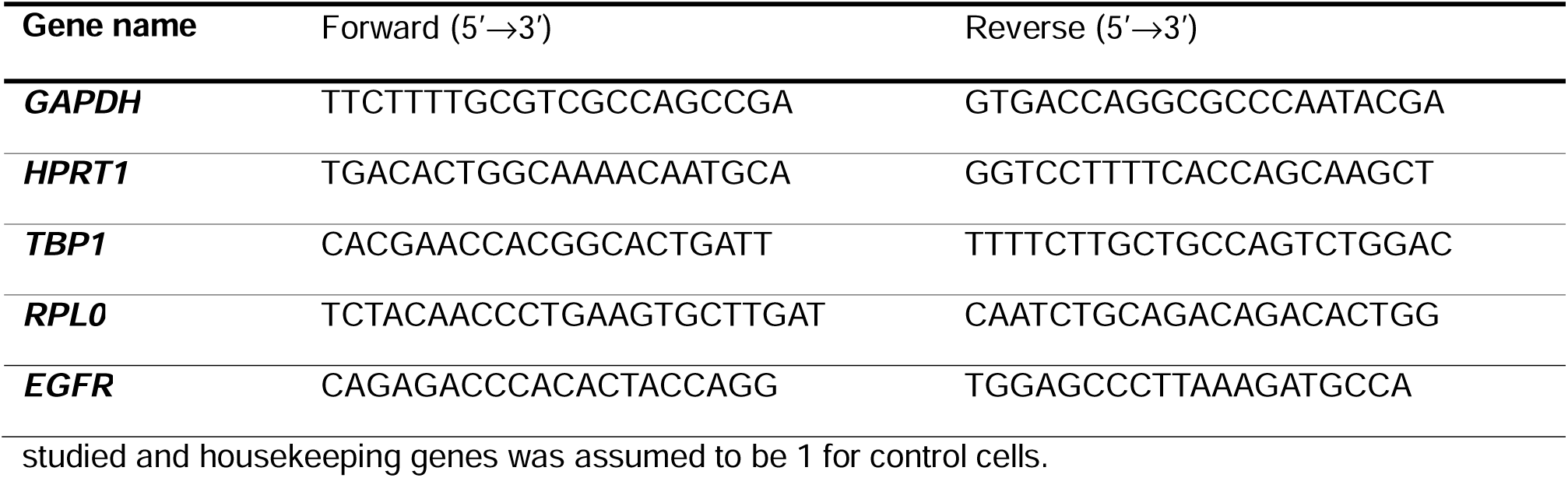
Gene primers.

### 2.16 Statistics

The Kolmogorov-Smirnov or Shapiro-Wilk test was applied to assess whether the data followed a normal distribution, while the homogeneity of variance was examined using Levene’s test. The results were expressed as mean ± standard deviation (SD) or standard error of the mean (SEM), depending on the context. One-way or two-way ANOVA followed by Dunnett’s was used for multiple comparisons of variation.

## 3. Results

### 3.1 Design and characterization of liposomes decorated with the R9 peptide

Two different strategies were used for the inclusion of R9 in the liposome formulation in order to have the R9 located on the surface of liposomes, which we described in our former study by Bellavita R. and co-authors [16]. The first strategy involved the binding of R9 to DSPE and the second to cholesterol, while having the peptide separated from anchor via PEG in both approaches. Such conjugates were purified and characterized by HPLC and mass spectroscopy.

NMR analysis shows that the reaction between the R9 peptide and DSPE-PEG_Maleimide was successful. In fact, the proton NMR spectrum (in Fig. S1) doesn’t show the typical signals of alkenes protons at ∼6.90 ppm from the intact maleimide, while typical signals of peptide, PEG, and DSPE moieties can be observed. The assignment is confirmed by 2D TOCSY spectrum. As regards R9_cholestreol the proton NMR spectrum (Fig. S2) shows NH amide resonances of arginine and lysine belonging to the moiety, whilst signals from cholesterol cannot unambiguously assigned. In fact, the typical cholesterol signal at 5.35 ppm (labeled by an asterisk *) is just detectable and that at 3.55 ppm overlaps to PEG resonances. However, a high field resonance at ca. 0.8 ppm may be assigned to a methyl group of cholesterol.

FTIR analysis was performed to support the identification of the R9-Chol construct by comparing the spectra of the R9 peptide, cholesterol, and the final conjugate. The main characteristic absorption bands related to the functional groups present in the individual components were evaluated.

In particular, the region between 3200 and 3400 cm⁻¹ was considered indicative of O-H and N-H stretching vibrations [23], while the region between 1100 and 1300 cm⁻¹ was analysed for bands associated with C-O and C-N stretching vibrations, in accordance with data reported in the literature for similar systems was considered supportive for the formation of the final construct [24].

A liposome composition based on DOPE (1,2-di-(9Z-octadecenoyl)-sn-glycero-3-phosphoethanolamine) and TAP (1,2-dimyristoyl-3-trimethylammonium-propane) are considered the best combination for siRNA delivery. TAP is a cationic lipid that provides the positive charges needed to ensure liposome internalization into the cell[25], while DOPE, helper lipid, with its two unsaturated fatty acid chains, exhibits fusogenic properties through its ability to generate a reversed hexagonal phase H^C^_II_, which destabilizes endosomal membranes, increasing the liposome’s ability to escape endosomally [26]. The DOPE:TAP lipid composition ratio in the liposome is approximately 1:2 as an alternative to the commonly used 1:1 ratio [27]. This change in ratio is due to the functionalization of the liposome with a peptide containing cholesterol in its structure [28].

In our assembly strategy, we first prepared liposomes with 20%, 30% and 40% in mol of peptide to compare hydrodynamic diameter and polydispersity index (PDI) by measuring dynamic light scattering, which correspond to liposome Z-average size and heterogeneity of the particle population, respectively. As shown in Table 3a and Table 3b, the size of liposomes fluctuated around 200 nm regardless of the peptide:lipid molecular ratio or the mode of peptide anchoring. Although the lack of clear shift in liposome size, the trend in polydispersity index became more pronounced as the R9-DSPE-PEG2000 content increased, indicating heterogeneous population or even unstable, aggregating system. In contrast, the opposite interdependence followed the increase in R9-CHOL content. Bearing in mind the desired size for liposomal/RNA delivery systems ranging from 50 to 200 nm and PDI from 0.1 to 0.2, which indicate well-controlled, likely monodisperse nanoparticle formulation, the most favourable parameters characterized the DOPE:TAP:R9-CHOL liposomes, when the components were combined in a molecular ratio of 2.1:5.6:2.3.

**Table 3a.** Characterization of uncomplexed liposomes made with R9-DSPE-PEG.

| Liposome composition | Molar ratio | Size (d. nm) | PDI | Zeta potential | Interaction with siRNA |
| --- | --- | --- | --- | --- | --- |
| DOPE:TAP:CHOL (Blank) | 7:2:1 | 172 ± 1 | 0.23 ± 0.02 | -27.9 mV ± 4.1 | - |
| DOPE: TAP:CHOL: R9-DSPE-PEG2000 | 6:1:1:2 | 152 ± 2 | 0.26 ± 0.01 | -0.7 mV ± 0.1 | - |
| DOPE:TAP:CHOL: R9-DSPE-PEG2000 | 4:2:1:3 | 249 ± 2 | 0.41 ± 0.02 | -3.9 mV ± 1.4 | - |
| DOPE:TAP:CHOL: R9-DSPE-PEG2000 | 3:2:1:4 | 195 ± 2 | 0.37 ± 0.01 | -3.9 mV ± 1.7 | - |

**Table 3b.** Characterization of uncomplexed liposomes made with R9-CHOL-PEG.

| Liposome composition | Molar ratio | Size (d. nm) | PDI | Zeta potential | Interaction with siRNA |
| --- | --- | --- | --- | --- | --- |
| DOPE:TAP:CHOL:R9-CHOL-PEG2 | 5:2:1:2 | 172 ± 14 | 0.31 ± 0.01 | --- (no peak) | - |
| DOPE:TAP:CHOL:R9-CHOL-PEG2 | 3:2:1:4 | 181 ± 4 | 0.24 ± 0.01 | +2.5 mV ± 0.8 | - |
| DOPE:TAP:CHOL:R9-CHOL-PEG2 | 2:3:1:4 | 246 ± 11 | 0.22 ± 0.01 | +2.3 mV ± 0.5 | - |
| DOPE:TAP:R9-CHOL-PEG2 | 2.1:5.6:2.3 | 160 ± 1 | 0.15 ± 0.07 | +18.5 mV ± 6.6 | + |

Since the complexation with siRNA requires cationic charges on the liposome surface, we measured the zeta potential, which indicates the effective surface charge. In case of DSPE-PEG2000-anchored nona-arginine peptide, the zeta potential was negative across all formulations. A plausible explanation for an unexpected reduction of surface charge is that the presence of PEG2000 partially hinders the R9. Moreover, the liposome surfaces carried a negative charge substantially limits the likelihood of siRNA binding, hence the liposomes with R9 anchored via DSPE-PEG2000 should be rejected as a potential carrier. This conclusion was confirmed by agarose gel electrophoresis, which indicated the lack of siRNA complexation regardless of DOPE: TAP:CHOL: R9-DSPE-PEG2000 molecular ratio (Fig. S3).

Whereas DSPE-PEG2000 failed to embed R9 peptide in DOPE:TAP:CHOL liposomes in a way to preserve positive charge on their surface, the relatively low R9 incorporation via CHOL-PEG2 was sufficient to shift the zeta potential toward positive values. However, the change in DOPE:TAP lipid ratio was not associated with further increase in zeta potential. Therefore, we decided to remove cholesterol from liposome bilayer and the resulted formulation DOPE:TAP:R9-CHOL-PEG2 in a molecular ratio of 2.1:5.6:2.3 achieved the highest z value of +18.5 mV ± 6.6. Reducing the amount of R9-chol peptide on the surface allowed the formation of liposomes with a smaller hydrodynamic diameter, maintaining their monodispersity at the PDI level of 0.15.

Concluding, among the tested formulations including two embedding options: DSPE-PEG2000 and CHOL-PEG2, only the latter composition emerged promising for siRNA complexation, because of desired size, relative monodispersity and cationic charge on their surface. The removal of cholesterol from lipid bilayer and its replacement with TAP substantially elevated surface cationic charge despite lower contribution of R9-CHOL-PEG2. The extended PEG chain negatively influenced liposome structural organization, but simultaneously masked the positive charge associated with nona-arginine (R9) peptide incorporation.

### 3.2 siRNA complexation with DOPE:TAP:R9-CHOL-PEG2 liposomes indicates optimal ratio of 1:77

Having the liposomal composition and component ratios chosen based on their size, polydispersity index and zeta potential, we investigated the complexation efficiency of siRNA with the liposomes. DOPE:TAP:R9-CHOL liposomes mixed at the ratio of 2.1:5.6:2.3 mol, which corresponds to 23% of R9-siRNA, were complexes with si*EGFR* in a wide molecular proportion ranging from 1:3.4 to 1:96 (Fig. 2A). By monitoring changes in rotational mobility of fluorescently labeled siRNA we observed a gradual increase in the siRNA fraction bound to liposome up to the molar ratio of 1:77, over which the binding equilibrium occurred. This siRNA:liposome saturation was confirmed by agarose gel electrophoresis, where a detectable free siRNA band was absent, thereby suggesting effective siRNA complexation (Fig. 2B).

**Fig. 1.**
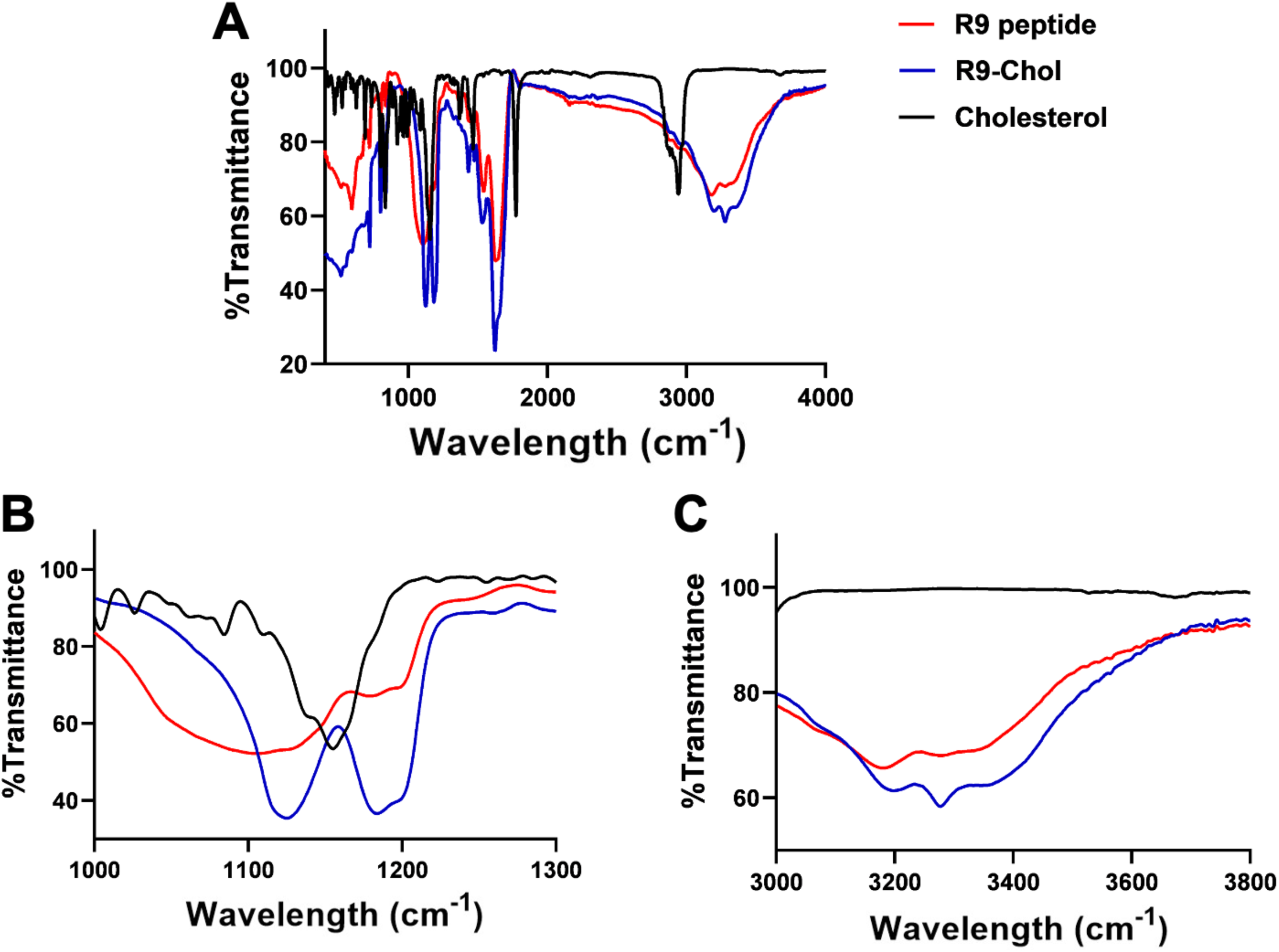
FTIR analysis of R9 peptide, cholesterol, and the R9-Chol. (A) FTIR spectra of the three samples. (B) Magnified view of the 1000-1300 cm⁻¹ region, highlighting spectral bands mainly associated with O-H and N-H stretching vibrations. (C) Magnified view of the 3000-3800 cm⁻¹ region, corresponding to C-O and C-N stretching vibrations.

**Fig. 2.**
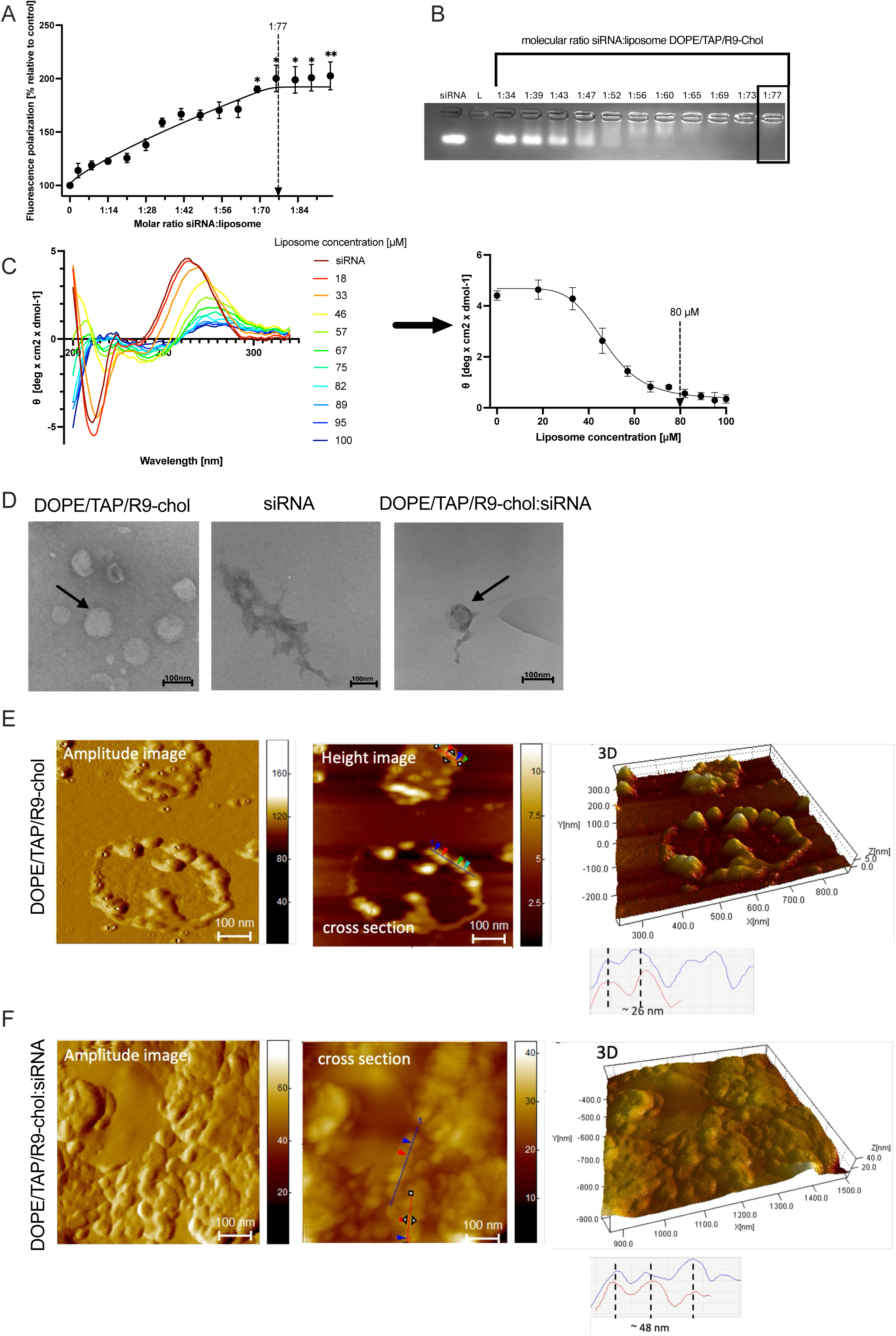
DOPE/TAP/R9-chol liposome interacts with siRNA. Analysis of the formation of the complex. (A) Change in fluorescence polarisation of siRNA-FITC under different molar ratios of siRNA:liposome by titration. Measurements performed in 10 mmol/L, pH = 7.4, 37°C phosphate buffer by PerkinElmer LS-50B spectrofluorometer. The control value (1:0) = 100% was the fluorescence polarisation of siRNA itself. The concentration of siRNA 0.1 μmol/L. The values are the mean ± SEM, n = 3. To compare multiple values, Shapiro-Wilk normality analysis and one-way analysis of variance followed by Dunnett’s post hoc test were performed. Statistical significance differences are marked * when p < 0.05, ** when p < 0.01 and *** when p < 0.001. (B) Agarose gel electrophoresis with an attempt to determine the most appropriate molar ratio between siRNA and DOPE/TAP/R9-Chol liposome (L). Samples containing 1 μmol/L siRNA per line and nanoparticles applied in the corresponding concentrations were prepared in PBS (phosphate-buffered saline; pH = 7.4) in the presence of GelRed stain. Gels were visualized upon transillumination at 525 nm. (C) Effect of DOPE/TAP/R9-Chol liposome on siRNA’s secondary structure (θ—ellipticity). Spectra CD of siRNA at the presence of liposome and changes in mean ellipticity at λ = 265 nm. Interpolation plotted based on the results. Measurements were conducted in λ = 200–320 nm in phosphate buffer, pH = 7.4. The concentration of siRNA 1 μmol/L. The values are the mean ± SEM, n = 3 Fig. (D) Transmission electron microscope images of the DOPE/TAP/R9-Chol liposome, siRNA and siRNA:liposome complex in molar ratio 1:77.

In order to check the possible conformational rearrangement of siRNA secondary structure upon complexation with DOPE:TAP:R9-CHOL liposomes, we made use of label-free circular dichroism (CD) spectroscopy (Fig. 2C) [29]. Liposome titration resulted in a decrease in siRNA ellipticity, which suggests that interaction with the liposomal carrier alters the structure of the siRNA duplex, likely due to electrostatic complexation. The maximal conformational rearrangement was achieved at the molecular ratio ∼1:80, which aligns well with results from fluorescence polarization (Fig. 2A) and gel electrophoresis (Fig. 2B), thereby also confirming that siRNA complexation by liposomes occurs at a molar ratio in the range of 1:70–1:80.

Comprehensive physicochemical characterization using transmission electron microscopy (TEM) (Fig. 2D) and atomic force microscopy (AFM) (Fig.2E) indicated the favourable morphology, nanoscale size distribution, and structural stability of the developed liposomal formulation, which are prerequisite for carrier for efficient siRNA delivery.

TEM analysis with negative staining was carried out to visualize the morphology of liposomes alone, naked siRNA, and siRNA–liposome complexes at a molar ratio of 1:77 (Fig.2D). Liposomes showed mostly spherical to quasi-spherical shapes with clear contours, typical of intact vesicular nanostructures. Against the electron-dense background, they appeared as bright vesicles surrounded by darker rims, matching the usual appearance of negatively stained lipid bilayers. The observed particle size mainly ranged between about 50 and 150 nm, though some larger vesicles and a few irregularly shaped particles were also seen. The lack of electron-dense internal structures further confirmed their vesicular nature. As expected for uncomplexed nucleic acids, siRNA appeared as dispersed filamentous and aggregated structures with irregular shapes and locally increased electron density. At the applied magnification, individual siRNA duplexes could not be resolved, so the observed structures likely represent higher-order aggregates or entangled bundles rather than single molecules. The morphology of siRNA–liposome complexes resembles a lipoplex-like arrangement, though definitive structural assignment is limited by the constraints of negative staining TEM.

The surface morphology of DOPE/TAP/R9-Chol liposomes and its alteration after complexation with siRNA was further examined by AFM. As typical for AFM imaging, liposomes underwent substantial deformation upon their contact with mica, leading to spontaneous formation of flattened, terrace-like spherical objects from 36 to 140 nm (Fig. 2E, S4). This heterogeneity may reflect differences in vesicle hydration and local surface interactions, possibly supported by reduced electrostatic repulsion at the liposome:mica interface. Following dilution to 100 µM, AFM imaging showed smaller bubble-like structures with diameters of about 26 nm (height image), while phase imaging indicated membrane thickness values consistent with lipid bilayer organization, in agreement with previous reports [30]. In the presence of siRNA, the system formed structures with an average high diameter of about 48 nm, thereby suggesting thickening of the liposome surface, most likely by siRNA complexation. Notably, no clear phase contrast was observed in this condition, also indicating altered mechanical or compositional surface properties of the resulting complexes (Fig. 2F, S4) [31].

In summary, analysis of biophysical parameters for siRNA interaction with DOPE/TAP/R9-Chol liposomes clearly indicates siRNA complexation on the liposome surface with optimal complexation molecular ratio of 1:77.

### 3.3 DOPE/TAP/R9-Chol:siRNA complexes demonstrate an improved zeta potential in physiologically-relevant buffer

Bearing in mind that physiological, *in vivo* conditions are characterized by the presence of ions and proteins, we tested the impact of phosphate buffer on size and surface charge of the liposome:siRNA complexes in the concentration range of 10-40 µM, which correspond to the siRNA concentration of 0.1-0.52 µM. Such siRNA concentrations are frequently used for gene silencing in cell culture [32], but also in *in vivo* studies [33].

As long as the liposome complexation with siRNA did not affect their size in water, the phosphate buffer alone caused considerable increase in liposome hydrodynamic parameter up to > 300 nm at their relatively low concentration (Fig. 3A). The same profile of changes was found for DOPE/TAP/R9-Chol:siRNA complexes suspended in the presence of ions. However, over the concentration of 20 µM their size gradually declined to <180 nm, which is acceptable for systemic delivery of nucleic acids. At the concentration of 38.5 µM, liposome alone and liposome:siRNA complexes preserved the desired low polydispersity (<0.3), which suggests colloidal stability under physiologically relevant ionic conditions (Fig. 3B).

**Fig. 3.**
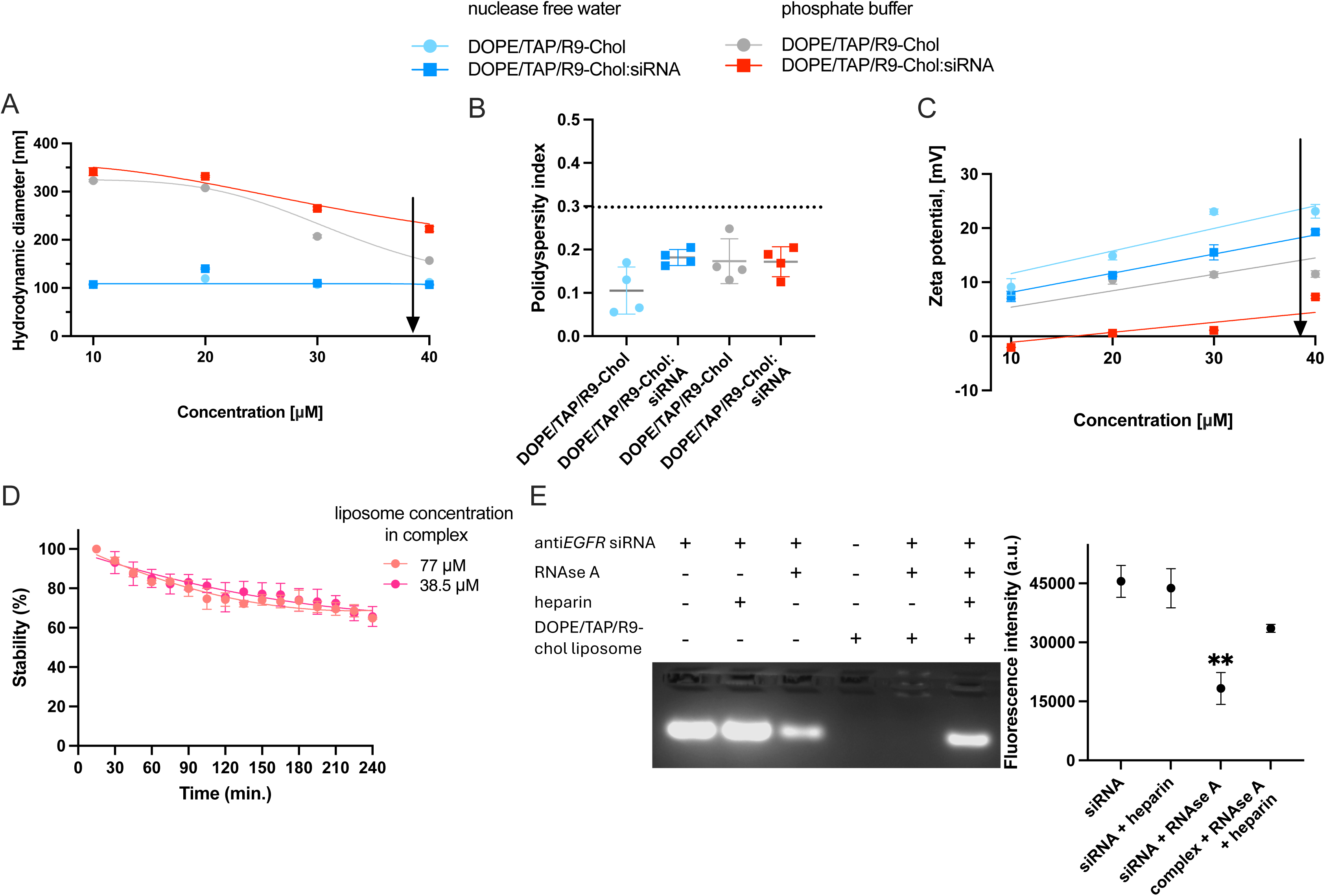
The size and charge of DOPE/TAP/R9-chol:siRNA remain constant in the desired range of siRNA. Measurement of the hydrodynamic diameter (A) with polydispersity index (B) of DOPE/TAP/R9-Chol liposomes and their complexes with si*EGFR* at different concentrations. The standard curve of interpolation is the change in concentration of the compound. Changes in the zeta potential (C) in different concentration of the formulation DOPE/TAP/R9-Chol alone and in complex with si*EGFR* at molar ratio 1:77 and in different solvents (nuclease free water and phosphate buffer, pH = 7.4). Determined simple linear regression. Changes in fluorescence polarization (D) of siRNA-FITC (0.1 μM) complexed with DOPE/TAP/R9-Chol liposome in two liposome concentration of 1:77 molar ratio in time, 37°C. The control value = 100% was the fluorescence polarization of complex after 15 min of incubation. The values are the mean ± SEM, n = 3. (E) Degradation of si*EGFR* uncomplexed and complexed with DOPE/TAP/R9-Chol liposome by RNAse A was visualized by agarose gel electrophoresis. and quantification of fluorescence intensity. DOPE/TAP/R9-Chol liposome concentration - 77 μM, siRNA - 1 μM, RNAse A - 1,25 μg/ml; complexes were formed in PBS solution in the presence of GelRed stain and siRNA digestion was performed at 37 C for 120 min. Release of siRNA from complexes after exposure to the enzyme was triggered by addition of 0.082 mg/ml heparin. Gels were visualized upon transillumination at 525 nm and fluorescence intensity was plotted in right panel. The difference between samples was tested using one-way ANOVA with Dunnett’s post hoc test, and statistically significant differences are marked with ** when p < 0.01.

Liposome complexation with siRNA declined their zeta potential and the difference remained constant regardless of the liposome concentration (Fig. 3C). Importantly, the shift to lower values was markedly stronger for liposomes and liposome complexes suspended in the phosphate buffer. DOPE/TAP/R9-Chol:siRNA complexes maintained relatively low zeta potential, which oscillated around 0 mV up 20 µM, and showed only a weak increase with further increasing complex concentration. The observed reduced profile of surface charge is particularly important for limiting unwanted interactions with serum proteins, which may otherwise promote aggregation, protein corona formation, and premature clearance. The observed processes are consistent with the Debye–Hückel theory [34].

Summing up, DOPE/TAP/R9-Chol:siRNA complexes maintained the expected particle size and narrow distribution in buffer, showing a more favorable zeta potential profile than in water, in line with better suitability for biological delivery uses.

### 3.4 DOPE/TAP/R9-chol liposomes form a stable complex with siRNA

The stability of the DOPE/TAP/R9-Chol:siRNA complexes was evaluated by examining both their temporal integrity under physiological-like conditions (Fig. 3D) and their capacity to protect siRNA from enzymatic degradation by ribonuclease A (Fig. 3E).

During the 4-hour observation period, chosen as a time window relevant for early systemic circulation and cellular delivery [35], the degree of complexation progressively declined over time, reaching approximately 70% at the end of the incubation, and did not vary between two concentrations studied (Fig. 3D). This led us to conclude that the storage of DOPE/TAP/R9-Chol:siRNA complexes may be associated with a reduced efficacy of siRNA delivery and that carefully calculated excess of siRNA may be required to achieve the desired biological effect [36].

In addition to complex stability, the protection of siRNA against nuclease-mediated degradation is a critical prerequisite for effective siRNA delivery, because these enzymes are abundant in the blood stream and inside cells. In Fig. 4E we show that siRNA complexation with DOPE/TAP/R9-Chol before incubation with RNAse A shields ribonucleic acid from the enzymatic attack, reducing accessibility of phosphodiester bonds to circulating ribonucleases. While using heparin for siRNA displacement from complexes, the protective effect of DOPE/TAP/R9-Chol liposomes was lost.

**Fig. 4.**
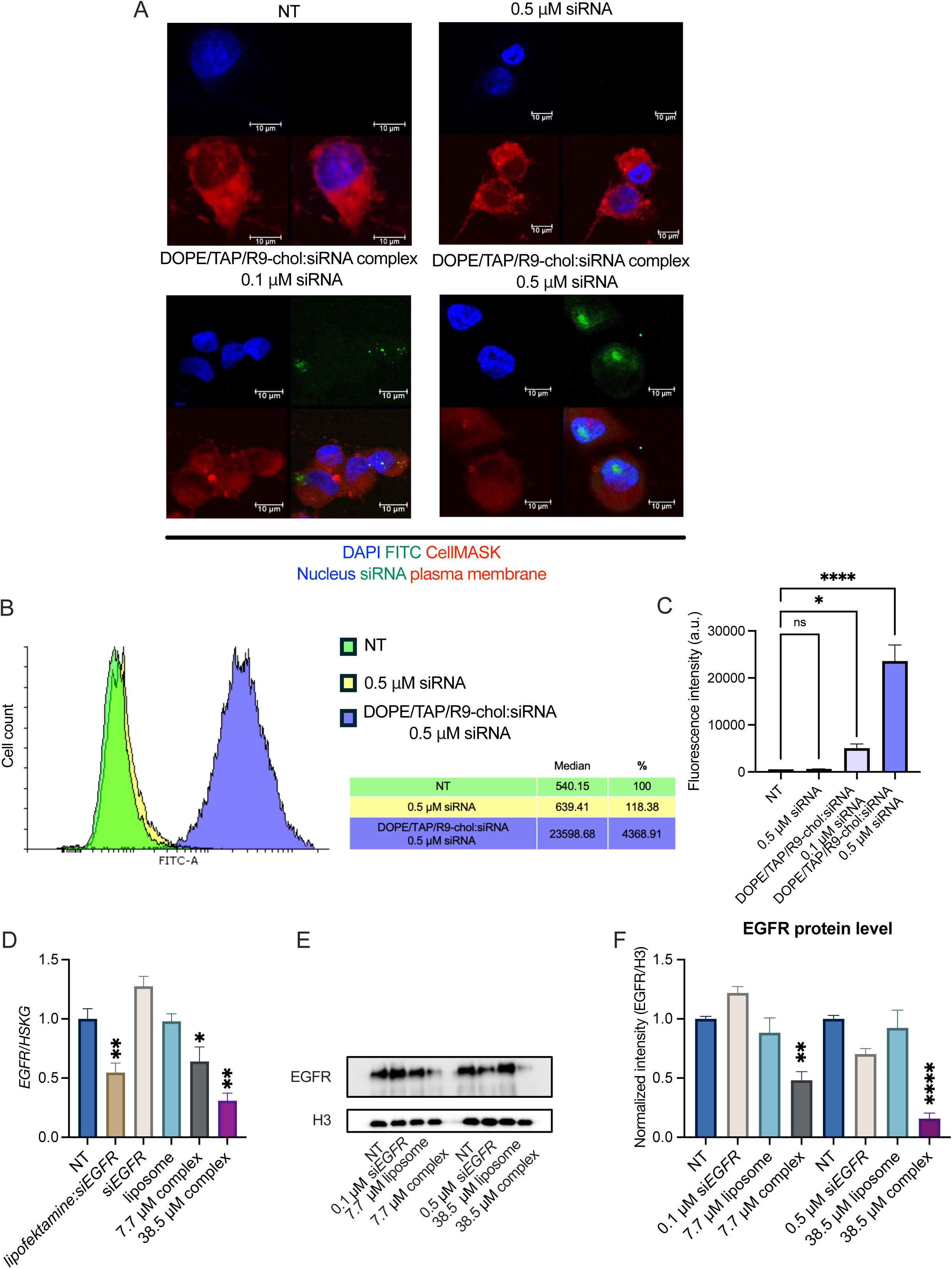
DOPE/TAP/R9-chol deliver siRNA into cells and reduce level of *EGFR* mRNA and expressed protein. (A) Example confocal microscopy images of si*EGFR* internalisation: MDA-MB-231 nontreated, treated with siRNA and siRNA:DOPE/TAP/R9-Chol complexes with liposome concentration of 7.7 µM and and 38.5 µM for 4 h. Complexes made at a molar concentration of 1:77. DNA was stained with DAPI, cell membrane stained with CellMASK. (LAS X, Leica Microsystems, Germany). DOPE:TAP:R9-Chol liposome complex increases siRNA accumulation in triple-negative breast cancer cells (B) Histogram shows the distribution of si*EGFR*-FITC fluorescence intensity quantified by flow cytometry. Each sample was tested in three biological replicates. Cells were treated with 0.5 µM siRNA and a 38.5 µM liposome complex, molar ratio of the complex 1:77 for 4h. The mean fluorescence of the cells and the fold change with respect to control cells (NT) is indicated in Table. (C) The mean fluorescence of cells treated with siRNA and two liposome complexes at concentrations of 7.7 and 38.5 µM, molar ratio of siRNA:liposome complex 1:77, which was read by flow cytometry. Results are presented as mean of median ± SD. Normality of data Shapiro-Wilk, and a one-way ANOVA followed by Dunnett’s post hoc test was performed for comparison of multiple samples. Samples were marked with * when p < 0.05, ** when p < 0.01, *** when p < 0.001 and ****p < 0.0001. (D) mRNA level was compared in cells 72h after their transfection with siCTRL, si*EGFR* by real-time PCR in MDA-MB-231. Silencing control performed using lipofectamine RNAiMAX:si*EGFR*. Raw values were normalized first to housekeeping genes (*TBP1*, *GAPDH*, *HPRT1* and *RPL0*), and then the ratio was assumed as 1 in control samples. The difference between samples was tested using one-way ANOVA with Dunnett’s post hoc test, and statistically significant differences are marked with * when p < 0.05 and ** when p < 0.01. (E) Comparison of Egfr protein level in MDA-MB-231 cells treated by liposome:si*EGFR* complex was studied by Western Blot. Histone H3 was used as a loading control. (F) Differential protein expression analysis by normalizes EGFR/H3 fold change. The difference between samples was tested using one-way ANOVA with Dunnett’s post hoc test, and statistically significant differences are marked with ** when p < 0.01 and **** p < 0.0001.

Concluding, our results show that the complexes underwent some time-dependent destabilization during incubation, but the liposomal formulation still retained a large portion of the complexed siRNA (∼70%) and successfully shielded the nucleic acid cargo from ribonuclease-mediated degradation.

### 3.5 Cellular uptake of DOPE/TAP/R9-Chol:si*EGFR* complex allows to efficiently silence *EGFR*

The presence of cell-penetrating cationic arginine-rich peptide on the liposome surface followed by the biophysical parameters of their complexes formed with siRNA were expected to promote efficient cellular internalization and intracellular accumulation of the transported siRNA [30]. To verify this hypothesis and monitor dose-dependent efficacy of siRNA delivery by the complex, we exposed MDA-MB-231 cells to two DOPE/TAP/R9-Chol:si*EGFR* concentrations: 7.7 µM and 38.5 µM, which correspond to 0.1 µM and 0.5 µM of siRNA, respectively (Fig. 3A).

Confocal microscopy analysis performed after 4 h cell incubation with DOPE/TAP/R9-Chol:siRNA-FITC complexes confirmed an intracellular siRNA-associated fluorescence signal. In cells exposed to lower dose of the complex, FITC-derived fluorescence was localized mostly in the cytoplasm, where canonical siRNA-mediated gene silencing occurs predominantly. At the higher complex concentration, si*EGFR*-FITC fluorescence was relatively uniformly distributed throughout the cytoplasm, with additional signal also observed within the nuclear region, which indicates very effective siRNA delivery [31][37][38]. As expected, siRNA alone did not enter cells and was not detected in cell cytoplasm.

To quantify more precisely the si*EGFR* accumulation inside cells across the entire cell population we used flow cytometry (Fig. 4B-C). In contrast to uncomplexed si*EGFR*, which failed to internalize, DOPE/TAP/R9-Chol:si*EGFR* complexes efficiently delivered siRNA into cells. This was observed by the relatively strong shift in the green, si*EGFR*-FITC-derived fluorescence. Importantly, a homogeneous distribution of the fluorescence intensity across the cell population suggested similar effectiveness of si*EGFR* delivery to most cells.

The concentration-dependent impact of DOPE/TAP/R9-Chol:si*EGFR* on mRNA level of *EGFR* we also observed after 48 h cell incubation with complexes (Fig. 5D). The studied transcript level was declined to 64% and to 31 % by 7.7 and 38.5 µM DOPE/TAP/R9-Chol:si*EGFR*, respectively. Importantly, the extent of *EGFR* silencing was comparable to lipofectamine RNAiMAX:si*EGFR* transfection, thereby suggesting that the developed formulation allows for effective intracellular siRNA delivery and leads to productive RNAi-mediated gene knock down.

To further validate the si*EGFR* delivery by DOPE/TAP/R9-Chol liposomes and gene silencing we tested the Egfr protein expression profile (Fig. 4E-F). As evidenced by western blot, Egfr level decreased substantially and in concentration-dependent manner 72 h after cell exposure to DOPE/TAP/R9-Chol:si*EGFR* complexes. Quantitative image analysis revealed a reduction in protein expression to approximately 48% and 15% of corresponding control levels for 7.7 µM and 38.5 µM complexes, respectively.

To conclude, the si*EGFR* internalization as well as *EGFR* silencing confirm that the developed liposomal formulation and its complexation with siRNA provides alternative for commercially available siRNA delivery platforms. The DOPE/TAP/R9-Chol:siRNA complexes efficiently overcame the cellular membrane barrier, enabling productive intracellular delivery of siRNA, which resulted in suppression of Egfr by reducing mRNA level.

## Discussion

The functionality and morphology of nanoparticles, including their shape, size and structure, play a key role in determining their properties and applications [39]. However, opportunities to improve existing nanoparticles to enhance their capabilities are being explored. The nona-arginine R9 peptide is known to be penetrating vie interacting with negative cell membranes, but also as carrier for siRNA and shRNA [40][41]. In our project, the above-mentioned cationic peptide was successfully anchored in the liposome lipid bilayer via cholesterol-PEG2, hence creating the efficient platform for siRNA delivery and gene silencing. Such a design of the DOPE/TAP/R9-Chol:siRNA platform offers a flexible foundation for further engineering aimed at targeted delivery and incorporation into lipid bilayer molecules targeting specific cancer markers. Such functionalization may include monoclonal antibodies, receptor-binding peptides, aptamers and many other, to significantly improve tumor selectivity. As we discussed in our recently published paper on *EGFR* targeting, high expression of this gene is associated with specific profile of other gene expression in triple-negative breast cancer, which can serve both as markers of Egfr-dependency and as cell-recognition elements for site-specific delivery [42].

The designed and developed DOPE/TAP/R9-Chol:siRNA lipoplexes display several physicochemical and functional features that are desirable for targeted nucleic acid delivery *in vitro* and *in vivo*. Their nanoscale size of approximate 180 nm at 38.5 µM, low polydispersity (<0.3), and stable colloidal behaviour in physiologically relevant ionic buffer are likely to support favourable biodistribution. These features are particularly important for drug delivery to tumors, where their capillary endothelium exhibits increased permeability, thereby facilitating the accumulation and prolonged retention of macromolecules that is known as EPR effect. Therefore, small, long-circulating liposomes are more likely to encounter these leaky capillaries and extravasate into the tumor microenvironment [43].

Importantly, the nearly neutral surface charge observed in buffered media may reduce non-specific interactions with serum proteins and help to minimize aggregation, protein corona formation, and rapid clearance by the mononuclear phagocyte system. The lack of positive charge on DOPE/TAP/R9-Chol:siRNA complexes at their concentration of 7.7 µM or relatively high hydrodynamic parameter (∼300 nm) did not compromise efficient intracellular siRNA delivery and gene silencing as evidenced by confocal microscopy, real-time PCR and western blot. Possibly, the reduced zeta potential likely reflects partial charge shielding in buffered ionic conditions, but DOPE/TAP/R9-Chol:siRNA complexes and, particularly, membrane-interacting R9 peptide preserved sufficient electrostatic character for interaction with cellular membrane. Moreover, the liposome bilayer components such as saturated phospholipid (DOPE) and cholesterol are known to decrease membrane fluidity, hence prolong the circulation time of liposomes [44]. The presence of TAP, a polycationic chemical structure, which prevents liposome-peptide-interaction, was shown to facilitate the intracellular release of nucleic acids [45]. Furthermore, the demonstrated protection of siRNA from ribonuclease-mediated degradation further supports suitability of DOPE/TAP/R9-Chol liposomes for systemic delivery of siRNA.

Complex stability is another critical parameter considered during the selection of suitable carriers for nucleic acid delivery. Even though we observed time-dependent destabilization as seen as a drop in complexation efficiency to about 70% after 4 h, the complexes stayed functionally active and kept their ability to deliver siRNA and trigger effective gene silencing at both mRNA and protein levels. Earlier studies provided experimental evidence that lipid-mediated siRNA internalization happens quickly, usually within minutes or during the two hours after exposure, while the first 4 h form a key period for intracellular uptake and cytosolic release required for effective RNA knock down [46][47]. These observations may also suggest that full structural stability might not be necessary, as long as the carrier keeps enough integrity throughout the biologically relevant delivery period.

Incorporation of the R9-Chol peptide into the DOPE/TAP liposomal formulation was intended to enhance siRNA binding through additional electrostatic interactions provided by the cationic arginine residues. Despite the incorporation of R9 into the lipid formulation via DSPE-PEG2000 anchoring, no positive surface charge was detected. A likely explanation of this phenomenon is steric shielding of the cationic R9 residues by the extended PEG2000 chains, which could mask the positive charges, or conformational back-folding of the PEG2000–R9 conjugate due to the high flexibility of PEG, which may lessen the electrostatic exposure of the arginine residue into the aqueous environment [48][49]. In line with these hypotheses, the use of a short PEG spacer in combination with cholesterol for R9 anchoring enabled the generation of a positive surface charge on the DOPE:TAP:CHOL:R9-CHOL liposomes at R9-CHOL-PEG molar ratio of 40%. However, the high content of free and R9-PEG2-conjugated cholesterol in the lipid bilayer prevented the generation of a sufficiently positive zeta potential required for efficient siRNA complexation [50]. Therefore, the composition of liposomes was limited to only two lipids: DOPE and TAP, which substantially increased the zeta potential and allowed for effective siRNA complexation.

Concluding, our study provides experimental evidence for the new siRNA delivery platform based on rationally designed R9-functionalized DOPE/TAP liposomes, which enables efficient siRNA complexation, protection, and effective intracellular accumulation. The developed system showed desirable physicochemical characteristics and effective suppression of *EGFR* expression *in vitro*. Altogether, these results suggest that this modular liposomal system could serve as a promising foundation for the continued development of targeted siRNA-based cancer therapies.

## Supporting information

whole WB images

Statistics

## Author contributions

Conceptualization: E.PW., M.B., S.G.; methodology: P.B., E.PW., AR; validation: P.B., S.B.; investigation: P.B. (DLS, CD, TEM, electrophoresis, fluorescence polarisation, flow cytometry, RT-PCR, western blot), S.B. (peptide synthesis and purification, DLS), T.M. (AFM), E.PW. (TEM), K.P. (liposome preparation), L.F. (NMR, FTIR); writing—original draft preparation: P.B and S.B.; writing — review and editing: E.PW., S.G. A.R., A.F., R.B.; supervision: A.R., E.PW., M.B., S.G.; project administration: E.PW., S.G.

## Data availability

The data used in this article have been deposited in the University of Lodz Repository. Raw data are available upon reasonable request to the corresponding author.

## Acknowledgements

The authors wish to thank Magdalena Gapińska, Łucja Balcerzak and Sława Glińska from the Laboratory of Microscopic Imaging & Specialized Biological Techniques, Faculty of Biology & Environmental Protection, University of Lodz, for their technical assistance.

International cooperation supported by joint research projects between Poland and Italy NAWA Canaletto entitled ‘Nanoparticles in the treatment of triple-negative breast cancer’.

---

**Scheme. 1.**
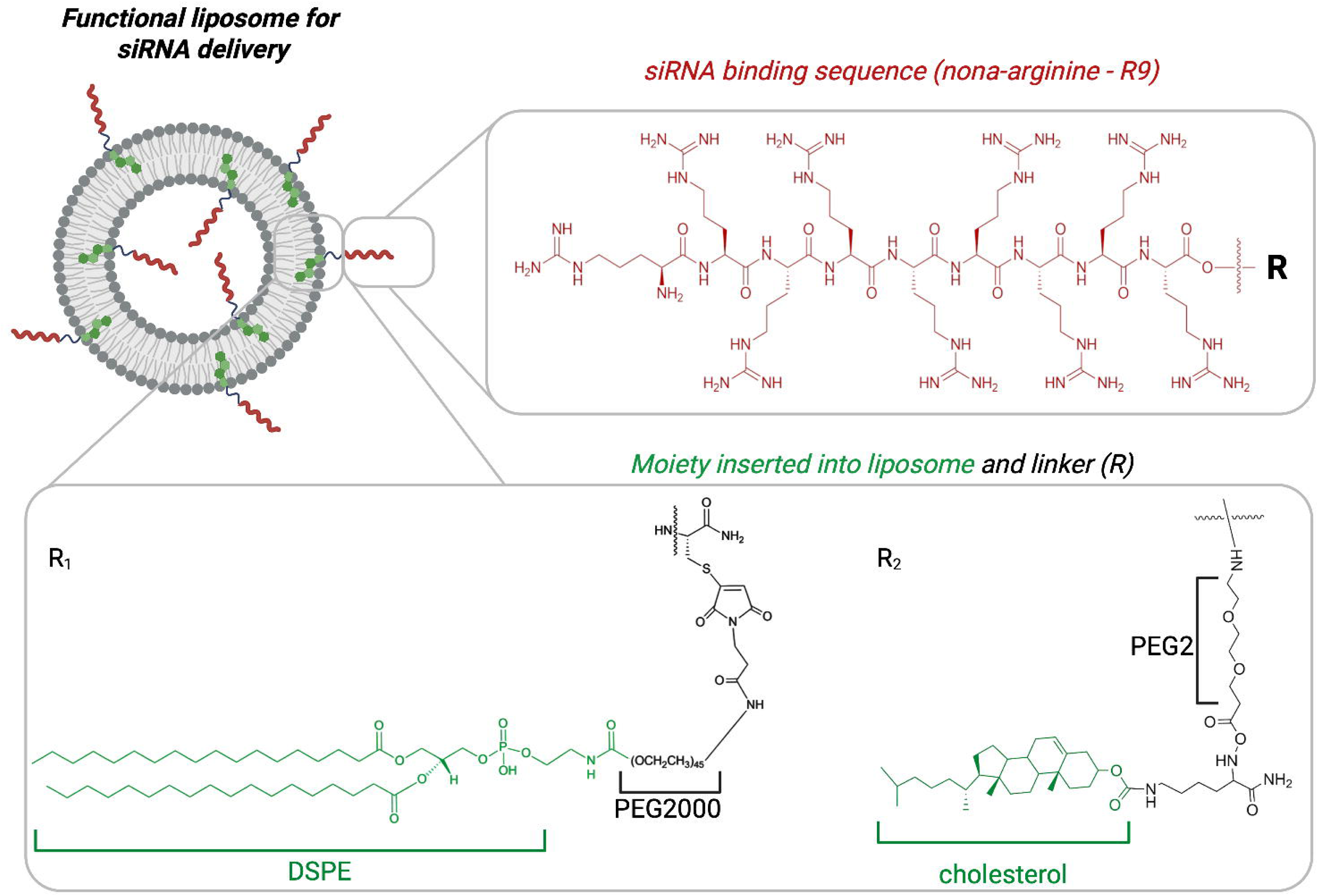
Suggested shape of DOPE/TAP/ liposome and structure of R9-DSPE-PEG2000 (R_1_) and R9-Chol (R_2_) peptides. Created in BioRender

## References

[1] H. Nsairat, D. Khater, U. Sayed, F. Odeh, A. Al Bawab, W. Alshaer, Liposomes: structure, composition, types, and clinical applications, Heliyon 8 (2022). 10.1016/j.heliyon.2022.e09394.

[2] Y. Jiang, W. Li, Z. Wang, J. Lu, Lipid-Based Nanotechnology: Liposome, Pharmaceutics 16 (2024). 10.3390/pharmaceutics16010034.

[3] P. Liu, G. Chen, J. Zhang, A Review of Liposomes as a Drug Delivery System: Current Status of Approved Products, Regulatory Environments, and Future Perspectives, Molecules 27 (2022) 1372. 10.3390/molecules27041372.

[4] E. Perillo, S. Porto, A. Falanga, S. Zappavigna, P. Stiuso, V. Tirino, V. Desiderio, G. Papaccio, M. Galdiero, A. Giordano, S. Galdiero, M. Caraglia, Liposome armed with herpes virus-derived gH625 peptide to overcome doxorubicin resistance in lung adenocarcinoma cell lines, Oncotarget 7 (2016) 4077–4092. 10.18632/oncotarget.6013.

[5] B.P. Meloni, F.L. Mastaglia, N.W. Knuckey, Cationic Arginine-Rich Peptides (CARPs): A Novel Class of Neuroprotective Agents With a Multimodal Mechanism of Action, Front. Neurol. 11 (2020). 10.3389/fneur.2020.00108.

[6] H. Yokoo, T. Misawa, T. Kato, M. Tanaka, Y. Demizu, M. Oba, Development of delivery carriers for plasmid DNA by conjugation of a helical template to oligoarginine, Bioorg. Med. Chem. 72 (2022) 116997. 10.1016/j.bmc.2022.116997.

[7] Y. Ando, H. Nakazawa, D. Miura, M. Otake, M. Umetsu, Enzymatic ligation of an antibody and arginine 9 peptide for efficient and cell-specific siRNA delivery, Sci. Rep. 11 (2021) 21882. 10.1038/s41598-021-01331-1.

[8] R. Anjous, P. Kavyashree, A. Saha, Strategies to improve intracellular delivery of arginine-rich cell-penetrating peptides, J. Mater. Chem. B 14 (2026) 3596–3619. 10.1039/D5TB02935J.

[9] A. Hibbitts, A.M. O’Connor, J. McCarthy, É.B. Forde, G. Hessman, C.M. O’Driscoll, S.-A. Cryan, M. Devocelle, Poly(ethylene glycol)-Based Peptidomimetic “PEGtide” of Oligo-Arginine Allows for Efficient siRNA Transfection and Gene Inhibition, ACS Omega 4 (2019) 10078–10088. 10.1021/acsomega.9b00265.

[10] Y. Choi, Kang, Park, Kang, Park, Folic acid-tethered Pep-1 peptide-conjugated liposomal nanocarrier for enhanced intracellular drug delivery to cancer cells: conformational characterization and in vitro cellular uptake evaluation, Int. J. Nanomedicine (2013) 1155. 10.2147/IJN.S39491.

[11] L.A. Orofiamma, D. Vural, C.N. Antonescu, Control of cell metabolism by the epidermal growth factor receptor, Biochim. Biophys. Acta Mol. Cell Res. 1869 (2022). 10.1016/j.bbamcr.2022.119359.

[12] J. Dancey, Targeting epidermal growth factor receptor—are we missing the mark?, 2003. 10.1016/S0140-6736(03)13810-X.

[13] R. Bellavita, A. Maione, S. Braccia, M. Sinoca, S. Galdiero, E. Galdiero, A. Falanga, Myxinidin-Derived Peptide against Biofilms Caused by Cystic Fibrosis Emerging Pathogens, Int. J. Mol. Sci. 24 (2023). 10.3390/ijms24043092.

[14] E. de Alteriis, A. Maione, A. Falanga, R. Bellavita, S. Galdiero, L. Albarano, M.M. Salvatore, E. Galdiero, M. Guida, Activity of Free and Liposome-Encapsulated Essential Oil from Lavandula angustifolia against Persister-Derived Biofilm of Candida auris, Antibiotics 11 (2021) 26. 10.3390/antibiotics11010026.

[15] T. Barra, A. Falanga, R. Bellavita, V. Laforgia, M. Prisco, S. Galdiero, S. Valiante, gH625-liposomes deliver PACAP through a dynamic in vitro model of the blood–brain barrier, Front. Physiol. 13 (2022). 10.3389/fphys.2022.932099.

[16] R. Bellavita, S. Braccia, M. Piccolo, P. Bialecki, M.G. Ferraro, S.F. Graziano, E. Esposito, F. Donadio, M. Bryszewska, C. Irace, E. Pedziwiatr-Werbicka, A. Falanga, S. Galdiero, Shielding siRNA by peptide-based nanofibers: An efficient approach for turning off EGFR gene in breast cancer, Int. J. Biol. Macromol. 292 (2025). 10.1016/j.ijbiomac.2024.139219.

[17] E. Pędziwiatr-Werbicka, M. Gorzkiewicz, S. Michlewska, M. Ionov, D. Shcharbin, B. Klajnert-Maculewicz, C.E. Peña-González, J. Sánchez-Nieves, R. Gómez, F.J. de la Mata, M. Bryszewska, Evaluation of dendronized gold nanoparticles as siRNAs carriers into cancer cells, J. Mol. Liq. 324 (2021) 114726. 10.1016/j.molliq.2020.114726.

[18] S. Zawadzki, Á. Martín-Serrano, E. Okła, M. Kędzierska, S. Garcia-Gallego, P.O. López, F.J. de la Mata, S. Michlewska, T. Makowski, M. Ionov, E. Pędziwiatr-Werbicka, M. Bryszewska, K. Miłowska, Synthesis and biophysical evaluation of carbosilane dendrimers as therapeutic siRNA carriers, Sci. Rep. 14 (2024). 10.1038/s41598-024-51238-w.

[19] K. Gronkowska, S. Michlewska, T. P³oszaj, M. Strachowska, A. Stępień, M. Borowiec, A. Bednarek, A. Robaszkiewicz, Targeting of Brahma-related gene-1 (BRG1) overcomes paclitaxel-induced multidrug resistance caused by overexpression of the subset of ATP-binding cassette (ABC) transporters, J. Pharmacol. Exp. Ther. 392 (2025) 103772. 10.1016/j.jpet.2025.103772.

[20] M. Strachowska, K. Gronkowska, M. Sobczak, M. Grodzicka, S. Michlewska, K. Kołacz, T. Sarkar, J. Korszun, M. Ionov, A. Robaszkiewicz, I-CBP112 declines overexpression of ATP-binding cassette transporters and sensitized drug-resistant MDA-MB-231 and A549 cell lines to chemotherapy drugs, Biomedicine & Pharmacotherapy 168 (2023) 115798. 10.1016/j.biopha.2023.115798.

[21] K. Gronkowska, S. Michlewska, A. Robaszkiewcz, Activity of Lysosomal ABCC3, ABCC5 and ABCC10 is Responsible for Lysosomal Sequestration of Doxorubicin and Paclitaxel-Oregongreen488 in Paclitaxel-Resistant Cancer Cell Lines, Cellular Physiology and Biochemistry 57 (2023) 360–378. 10.33594/000000663.

[22] K. Gronkowska, K. Kołacz-Milewska, S. Michlewska, T. Płoszaj, M. Borowiec, A. Robaszkiewicz, HIF1A, BRG1, and p300 interaction confers paclitaxel-induced drug resistance by enabling the overexpression of ABCC genes, Molecular Therapy Oncology 33 (2025) 201049. 10.1016/j.omton.2025.201049.

[23] A. Arrout, Y. El Ghallab, A. Hirri, R. Aït Mouss, I. Yamari, M. Rachid Lefriyekh, A. Elmakssoudi, A. Ait Haj Said, Prediction of cholesterol content in gallstones by FTIR spectroscopy coupled with chemometric tools, Microchemical Journal 199 (2024) 109956. 10.1016/j.microc.2024.109956.

[24] V. Bali, Y. Khajuria, V. Manyar, P.K. Rai, U. Kumar, C. Ghany, S. Tripathi, V.K. Singh, Elemental studies and mapping of cholesterol and pigment gallstones using scanning electron microscopy–energy dispersive spectroscopy, X-Ray Spectrometry 53 (2024) 487–498. 10.1002/xrs.3403.

[25] B.-K. Kim, G.-B. Hwang, Y.-B. Seu, J.-S. Choi, K.S. Jin, K.-O. Doh, DOTAP/DOPE ratio and cell type determine transfection efficiency with DOTAP-liposomes, Biochimica et Biophysica Acta (BBA) - Biomembranes 1848 (2015) 1996–2001. 10.1016/j.bbamem.2015.06.020.

[26] V. Vysochinskaya, S. Shishlyannikov, Y. Zabrodskaya, E. Shmendel, S. Klotchenko, O. Dobrovolskaya, N. Gavrilova, D. Makarova, M. Plotnikova, E. Elpaeva, A. Gorshkov, D. Moshkoff, M. Maslov, A. Vasin, Influence of Lipid Composition of Cationic Liposomes 2X3-DOPE on mRNA Delivery into Eukaryotic Cells, Pharmaceutics 15 (2022) 8. 10.3390/pharmaceutics15010008.

[27] Y. Zhang, H. Li, J. Sun, J. Gao, W. Liu, B. Li, Y. Guo, J. Chen, DC-Chol/DOPE cationic liposomes: A comparative study of the influence factors on plasmid pDNA and siRNA gene delivery, Int. J. Pharm. 390 (2010) 198–207. 10.1016/j.ijpharm.2010.01.035.

[28] A. Lechanteur, V. Sanna, A. Duchemin, B. Evrard, D. Mottet, G. Piel, Cationic Liposomes Carrying siRNA: Impact of Lipid Composition on Physicochemical Properties, Cytotoxicity and Endosomal Escape, Nanomaterials 8 (2018) 270. 10.3390/nano8050270.

[29] M. Danaei, M. Dehghankhold, S. Ataei, F. Hasanzadeh Davarani, R. Javanmard, A. Dokhani, S. Khorasani, M.R. Mozafari, Impact of Particle Size and Polydispersity Index on the Clinical Applications of Lipidic Nanocarrier Systems, Pharmaceutics 10 (2018) 57. 10.3390/pharmaceutics10020057.

[30] A. Botet-Carreras, M.B. Marimon, R. Millan-Solsona, E. Aubets, C.J. Ciudad, V. Noé, M.T. Montero, Ò. Domènech, J.H. Borrell, On the uptake of cationic liposomes by cells: From changes in elasticity to internalization, Colloids Surf. B Biointerfaces 221 (2023) 112968. 10.1016/j.colsurfb.2022.112968.

[31] N. Li, H. Yang, Z. Yu, Y. Li, W. Pan, H. Wang, B. Tang, Nuclear-targeted siRNA delivery for long-term gene silencing, Chem. Sci. 8 (2017) 2816–2822. 10.1039/C6SC04293G.

[32] L. Salim, C. McKim, J.-P. Desaulniers, Effective carrier-free gene-silencing activity of cholesterol-modified siRNAs, RSC Adv. 8 (2018) 22963–22966. 10.1039/C8RA03908A.

[33] M. Jayaraman, S.M. Ansell, B.L. Mui, Y.K. Tam, J. Chen, X. Du, D. Butler, L. Eltepu, S. Matsuda, J.K. Narayanannair, K.G. Rajeev, I.M. Hafez, A. Akinc, M.A. Maier, M.A. Tracy, P.R. Cullis, T.D. Madden, M. Manoharan, M.J. Hope, Maximizing the Potency of siRNA Lipid Nanoparticles for Hepatic Gene Silencing In Vivo**, Angew. Chem. Int. Ed. 51 (2012) 8529–8533. 10.1002/anie.201203263.

[34] L. Zhang, P. Winkler, Debye–Hückel screening and fluctuations, Chem. Phys. 329 (2006) 338–342. 10.1016/j.chemphys.2006.07.031.

[35] X. Cao, Y. Sun, P. Lu, M. Zhao, Fluorescence imaging of intracellular nucleases—A review, Anal. Chim. Acta 1137 (2020) 225–237. 10.1016/j.aca.2020.08.013.

[36] C. Safinya, Structures of lipid–DNA complexes: supramolecular assembly and gene delivery, Curr. Opin. Struct. Biol. 11 (2001) 440–448. 10.1016/S0959-440X(00)00230-X.

[37] S. Guang, A.F. Bochner, D.M. Pavelec, K.B. Burkhart, S. Harding, J. Lachowiec, S. Kennedy, An Argonaute Transports siRNAs from the Cytoplasm to the Nucleus, Science (1979). 321 (2008) 537–541. 10.1126/science.1157647.

[38] S.Y. Berezhna, L. Supekova, F. Supek, P.G. Schultz, A.A. Deniz, siRNA in human cells selectively localizes to target RNA sites, Proceedings of the National Academy of Sciences 103 (2006) 7682–7687. 10.1073/pnas.0600148103.

[39] M. Honciuc, A. Honciuc, Morphological Design and Synthesis of Nanoparticles, Nanomaterials 14 (2024) 360. 10.3390/nano14040360.

[40] R. Taghizadeh Pirposhteh, E. Arefian, A. Arashkia, N. Mohajel, Nona-Arginine Mediated Anti-E6 ShRNA Delivery Suppresses the Growth of Hela Cells in vitro, Iran. Biomed. J. 27 (2023) 349–356. 10.61186/ibj.3963.

[41] K. Melikov, A. Hara, K. Yamoah, E. Zaitseva, E. Zaitsev, L.V. Chernomordik, Efficient entry of cell-penetrating peptide nona-arginine into adherent cells involves a transient increase in intracellular calcium, Biochemical Journal 471 (2015) 221–230. 10.1042/BJ20150272.

[42] P. Białecki, M. Adamiak, E. Pędziwiatr-Werbicka, A. Robaszkiewicz, Perspectives of EGFR co-targeting with nanocarriers for anti-TNBC purposes, Genes Dis. (2026) 102026. 10.1016/j.gendis.2025.102026.

[43] M.S. Gatto, M.P. Johnson, W. Najahi-Missaoui, Targeted Liposomal Drug Delivery: Overview of the Current Applications and Challenges,. 14 (2024) 672. 10.3390/life14060672.

[44] A. Nagayasu, K. Uchiyama, H. Kiwada, The size of liposomes: a factor which affects their targeting efficiency to tumors and therapeutic activity of liposomal antitumor drugs, Adv. Drug Deliv. Rev. 40 (1999) 75–87. 10.1016/S0169-409X(99)00041-1.

[45] D.A. Balazs, WT. Godbey, Liposomes for Use in Gene Delivery, J. Drug Deliv. 2011 (2011) 1–12. 10.1155/2011/326497.

[46] A. Robaszkiewicz, E. Wiśnik, Z. Regdon, K. Chmielewska, L. Virág, PARP1 facilitates EP300 recruitment to the promoters of the subset of RBL2-dependent genes, Biochimica et Biophysica Acta (BBA) - Gene Regulatory Mechanisms 1861 (2018) 41–53. 10.1016/j.bbagrm.2017.12.001.

[47] M. Hirsch, M. Helm, Live cell imaging of duplex siRNA intracellular trafficking, Nucleic Acids Res. 43 (2015) 4650–4660. 10.1093/nar/gkv307.

[48] W. Yang, T. Miyazaki, P. Chen, T. Hong, M. Naito, Y. Miyahara, A. Matsumoto, K. Kataoka, K. Miyata, H. Cabral, Block catiomer with flexible cationic segment enhances complexation with siRNA and the delivery performance in vitro, Sci. Technol. Adv. Mater. 22 (2021) 850–863. 10.1080/14686996.2021.1976055.

[49] X. Ge, L. Chen, B. Zhao, W. Yuan, Rationale and Application of PEGylated Lipid-Based System for Advanced Target Delivery of siRNA, Front. Pharmacol. 11 (2021). 10.3389/fphar.2020.598175.

[50] N. Pavlovic, J. Mijalković, B. Balanč, N. Luković, Z. Knežević-Jugović, Role of Cholesterol in Modifying the Physical and Stability Properties of Liposomes and In Vitro Release of VitaminB12, in: The IX International Congress “Engineering, Environment and Materials in Process Industry”—EEM2025, MDPI, Basel Switzerland, 2025: p. 10. 10.3390/engproc2025099010.

