## Supplementary Information for "Lipid anchor engineering controls cell-penetrating arginine-rich peptide presentation for efficient siEGFR liposomal delivery to triple-negative breast cancer cells"

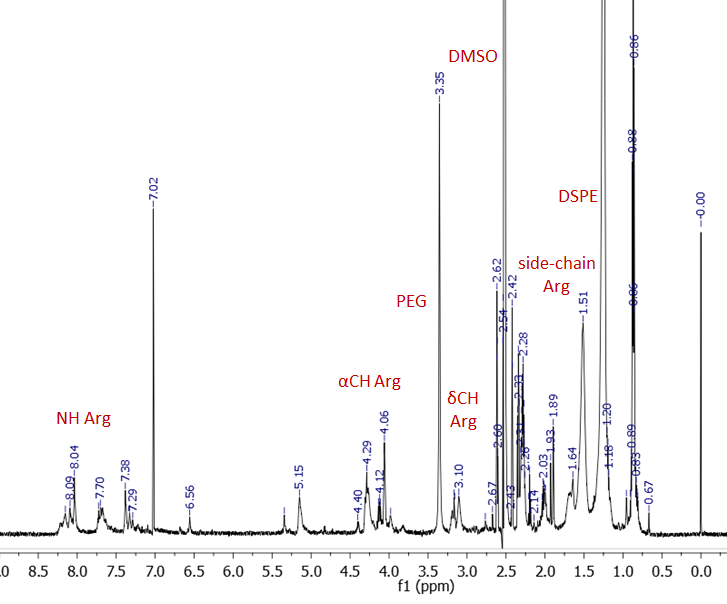

Fig. S1 NMR (SM) Proton NMR spectrum of R9_PEG_DSPE in DMSO at 298K and 700 MHz.

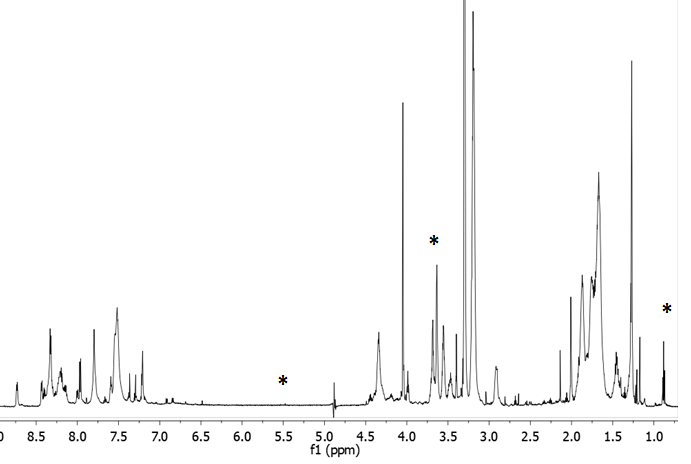

Fig. S2 NMR (SM) Proton NMR spectrum of R9_cholesterol in CD_3_OH at 298K and 600 MHz.

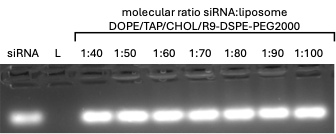

Fig. S3 Agarose gel electrophoresis with an attempt to determine the most appropriate molar ratio between siRNA and DOPE/TAP/R9-DSPE-PEG2000 liposome (L). Samples containing 1 μmol/L siRNA per line and nanoparticles applied in the corresponding concentrations were prepared in PBS (phosphate-buffered saline; pH = 7.4) in the presence of GelRed stain. Gels were visualized upon transillumination at 525 nm.

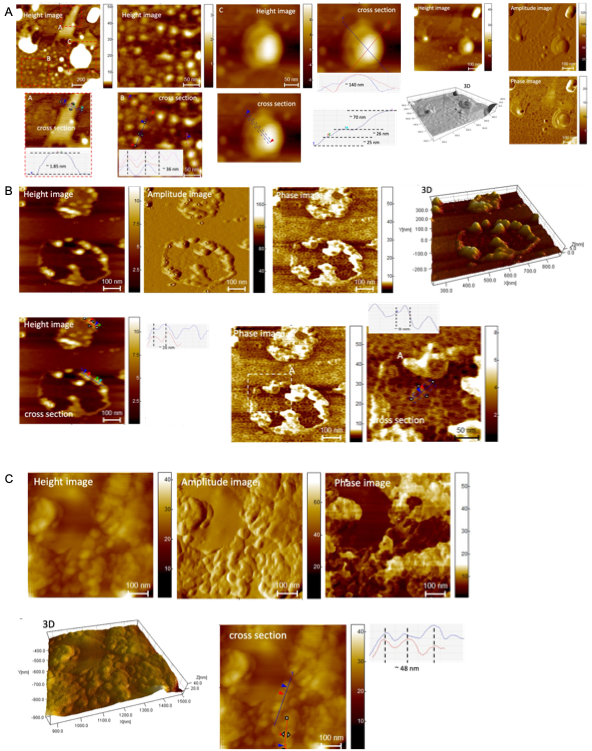

Fig. S4 Whole AFM images of DOPE/TAP/R9-chol liposomes and 3D visualization of its structure in 2 concentrations, (A) 1mM, (B) 100µM and (C) anti*EGFR* siRNA:liposome complex in a molar ratio of 1:77 (the optimal siRNA:liposome ratio) and 3D visualization of its structure).
