## Supplementary figures and images for "Lipid anchor engineering controls cell-penetrating arginine-rich peptide presentation for efficient siEGFR liposomal delivery to triple-negative breast cancer cells"

### whole WB images

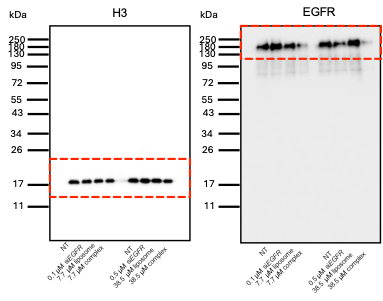

Figure S1 Whole WB Images
